# Compressive force from the nuclear envelope in dividing *S. pombe* alters binding of mitotic spindle proteins and regulates microtubule dynamics

**DOI:** 10.64898/2026.01.15.699759

**Authors:** Taylor Mahoney, Christopher Needham, Reem Hakeem, Marcus Begley, Christian Pagán Medina, Mary Williard Elting

**Affiliations:** North Carolina State University, Raleigh, NC; West Chester University of Pennsylvania, West Chester, PA; Rutgers University, Piscataway, NJ; Cluster for Quantitative and Computational Developmental Biology, Integrative Sciences Initiative, North Carolina State University, Raleigh, NC

## Abstract

During closed mitosis in *S. pombe*, the nuclear envelope and mitotic spindle must work together to ensure successful nuclear division. Previous work has demonstrated that mechanical force from the nuclear envelope, transmitted through spindle pole bodies, can reshape the spindle. However, it remains unclear how force from the nuclear envelope might regulate binding or other biochemical properties of microtubule associated proteins (MAPs) in the spindle. Here, we investigate how force reprograms the spindle with two approaches: chronically increasing nuclear envelope tension via the lipid synthesis inhibitor cerulenin, and acutely applying force to the nucleus through an optical trap. Both perturbations slow spindle elongation dynamics and reduce microtubule density. Despite this reduction, key spindle proteins Ase1 and Klp5 increase their density at the spindle midzone, indicating inward force from the nuclear envelope can alter MAP binding in the spindle. We find that while motor proteins Klp5 and Klp6 only minimally affect the spindle’s response to increased nuclear envelope force, the combination of removing Ase1 and increasing nuclear envelope force together rescues spindle microtubule stability. Together, our findings reveal that nuclear force on the spindle does not merely alter its shape, but is key in regulating its biochemistry to maintain force balance and ensure cellular function.

**SIGNIFICANCE:** Accurate chromosome segregation depends on the mechanical integrity of the mitotic spindle, yet how cells sense and respond to spindle destabilization remains poorly understood. Here, we show that disrupting the force balance between the nuclear envelope and spindle in the fission yeast *S. pombe* activates a compensatory response that promotes microtubule formation and restores spindle elongation. Through both optical trapping and pharmacological perturbation, we identify Ase1 as a critical stabilizer of spindle mechanics and demonstrate that cells can adapt to mechanical failure through regulation of spindle protein binding. These findings uncover a mechanotransduction feedback mechanism that links spindle force balance to cytoskeletal remodeling during mitosis.

## INTRODUCTION

In the mitotic spindle, force from elongating microtubules segregates chromosomes, ensuring that they are physically separated into two new daughter cells. Bundle elongation may require both sliding of microtubule motors and growth at microtubule plus-ends [1], [2], [3], while inward force can be generated by sliding or by microtubule depolymerization [2], [4]. These dynamics together exert a net force on chromosomes that delivers them to opposite ends of the cell [5]. In the case of closed mitosis, the spindle must further overcome compressive force from the nucleus itself [6], [7], [8]. This compressive force is concentrated at the spindle pole bodies (SPBs) [9], which embed in the nuclear envelope during mitosis [10], ensuring that extensile force from the spindle not only segregates chromosomes, but also reshapes the intact nucleus while doing so. Thus, closed mitosis requires coordination of physical structures with the biochemical processes that regulate them [11], [12].

The fission yeast *S. pombe* is an ideal system for disentangling these coordinated activities. Work over previous decades has established a basic framework of the process of spindle assembly and chromosome segregation in this system. The *S. pombe* mitotic spindle consists of 10-20 individual microtubules joined together in a single bipolar bundle [13], [14], [15], [16], [17]. Spindle elongation relies on extensile sliding by molecular motors such as Klp9 and Cut7 [3], while depolymerizing motors such as Klp5 and Klp6 help regulate microtubule growth and shrinkage [2], [14], and crosslinkers such as Ase1 ensure mechanical robustness of the bundle as a whole [17], [18], [19]. Together, this machinery powers the spindle as it rapidly elongates from nanometer to micron scale over the course of about half an hour, with length and timing that are highly consistent from cell to cell [18], [20]. The simple and stereotyped pattern of cell division and morphology has made the *S. pombe* spindle apparatus particularly amenable for mechanical perturbation and modeling [9], [21], [22], [23], [24], [25], [26]. However, the physical coordination between the mitotic spindle and the nuclear envelope is less completely understood.

During *S. pombe* mitosis, the nuclear envelope expands and undergoes extensive remodeling to accommodate the elongating spindle while initially remaining intact [27], [28]. Ultimately, the nuclear envelope must be topologically altered to enclose two daughter nuclei, and recent work has provided some insight into how this change is accomplished through local disassembly of nuclear pores in the narrow bridge [11], [12], [29], [30], though questions remain on how it coordinates with the spindle during this process. Pharmacological limitation of nuclear envelope expansion via the small molecule inhibitor cerulenin perturbs spindle elongation, often leading to bent spindles and malformed nuclei [6], [7], [8]. While the direct mechanical linkage that SPBs provide between the nuclear envelope and the mitotic spindle [13], [31] may cause this response, less direct explanations are also possible. For example, chromosome entanglements induced by cycles of osmotic shock also induce spindle bending, likely due to compressive forces transmitted between SPBs by chromosomes rather than the nuclear envelope [32], suggesting that lagging chromosomes, produced by such entanglements or other means, can also alter spindle morphology. Thus, more work is needed to understand how the spindle and the nuclear envelope effectively function together.

To ensure spindle function that accurately delivers chromosomes in opposition to compressive force, spindle microtubule bundles likely require effective crosslinking and appropriate microtubule length regulation. Ase1 is considered the primary antiparallel microtubule crosslinker in *S. pombe* [21], [33]. It is essential for both spindle stability and proper spindle elongation; previous work demonstrates that its absence causes disruptions to spindle elongation [15], [16], [17], [18], [33], [34]. In terms of regulating microtubule length, the plus-end-directed molecular motors Klp5 and Klp6 are key [35]. These two motors heterodimerize, and together promote depolymerization at microtubule ends which helps regulate spindle length and is crucial for normal chromosome movement [36], [37]. There is also some evidence that Klp5, potentially because of its ability to homodimerize [2], [35], may contribute to spindle bundling, as its loss results in occasional tri-polar spindles [2], [35].

Disentangling mechanical response from biochemical signaling cascades remains a key challenge to a complete physical picture of closed mitosis. Here, we address this knowledge gap, by applying two complementary approaches that both increase inward force on the spindle: disrupting membrane synthesis pharmacologically and pushing on the nuclear envelope with an optical trap. We expect both of these perturbations to increase nuclear envelope tension, and thereby to cause inward force on the spindle exerted through the SPBs. The former approach provides a sustained and systemic perturbation to the mechanics of the nucleus as a whole, whereas the second allows us to exert force that is more targeted in space and time. Spindles exhibit strikingly similar responses to both perturbations, suggesting that the effects we detect are indeed mechanically mediated. We uncover evidence that mechanical force exerted through the nuclear envelope induces changes in the molecular regulation of the spindle, suggesting how two critical structures in closed cell division coordinate their physical rearrangements.

## MATERIALS AND METHODS

### Cell culture and imaging

#### Fission yeast strains and culture

For details of *S. pombe* strains used in this work, see Table 1. We performed new crosses by tetrad dissection using standard methods [38]. We cultured all strains at 25 °C on YE5S plates using standard techniques [38]. For imaging, we grew liquid cultures in YE5S media at 25 °C with shaking by a rotating drum for 12-24 hours before imaging. To ensure that cells were in growth phase for imaging, we measured OD595 with a target range of 0.1-0.2. If cells had grown beyond this point, we diluted them and allowed them to recover for ~1 hour before imaging.

**Table 1.** *S. pombe* Strains.

| Genotype | Original source | Strain identifier |
| --- | --- | --- |
| GFP-atb2:kanMX, ade6-, leu1-32, ura4-D18, h+ | Gift of F. Chang (FC2861) | MWE2 |
| ase1::KanMX leu1-32::SV40-GFP-atb2[LEU1], leu1-32, ura4-D18, his7+, h- | Gift of F. Chang (FC1984) [19] | MWE3 |
| nmt41-GFP-atb2-hpnMX6, Cut11-meGFP-KanMX6 | Gift of C. Laplante (CL579-2) | MWE37 |
| ade6-216, leu1-32, lys1-131, ura4-D18, ase1::ase1-GFP-HA-Kanr, h90 | NBRP (FY14688) [40] | MWE60 |
| klp6::his3+, ade6-M210, his-D1, leu1-32, ura4-D18 | Gift of M. Betterton/J. Richard McIntosh (McI386) [37] | MWE73 |
| klp5::ura4+, nmt41-GFP-atb2-hphMX6, Cut11-meGFP-KanMX6 | This study (cross of MWE141 and MWE37) | MWE75 |
| klp6::his3+, GFP-atb2:kanMX | This study (cross of MWE73 and MWE2) | MWE76 |
| his5+:ptdh1:NLSSV40-sfGFP-NLSSV40:natMX | NBRP (FY38490) | MWE116 |
| klp5::ura4+, ade6-m216, his3-D1, leu1-32, ura4-D18, h+ | Gift of M. Betterton/J. Richard McIntosh (McI381) [37] | MWE141 |
| klp5+:GFP:Ura4+, ade6-M210, his3-D1, leu1-32, ura4-D18 | Gift of M. Betterton/J. Richard McIntosh (McI485) [37] | MWE142 |

As a method of increasing nuclear envelope tension in *S. pombe*, we treated cells with 0.3 mM cerulenin (from stock solutions at 50 mM in DMSO) for a period of 1 to 3 hours (depending on the batch of stock, see below) before imaging, as previously described [39]. For each new batch of cerulenin, we determined treatment time by screening for the first consistent appearance of curved spindles.

#### Imaging sample preparation

We prepared live samples for imaging as previously described [6], [25]. Briefly, prior to imaging, we placed yeast cells onto a gelatin pad (250 mg/mL in YE5S) that was adhered to a microscope glass slide. We first heated a gelatin solution in a tabletop dry heat bath at 90°C just until molten and then pipetted a 10 µL drop onto a slide, flattened the drop with a coverslip, and waited a minimum of 15 min for the gelatin pad to solidify. We then spun down a 1 mL volume of cells (previously grown in YE5S liquid growth media overnight and diluted to an OD of 0.1-0.2) using a tabletop centrifuge. We decanted as much of the supernatant as possible without disturbing the pellet and resuspended the cells in the remaining supernatant. Next, we pipetted 2 µL of the cell suspension onto the center of the gelatin pad, placed the coverslip on top of the drop, and sealed the edges of the coverslip using VALAP (1:1:1, Vaseline:lanolin:paraffin). We imaged all samples at room temperature (≈ 22°C).

#### Live cell spinning disc confocal imaging and laser ablation

We performed spinning disk confocal live imaging and laser ablation experiments as described previously [6], [25], [41]. We captured live videos using a Nikon Ti-E stand on an Andor Dragonfly spinning disk confocal fluorescence microscope containing a spinning disk dichroic Chroma ZT405/488/561/640rpc, 488 nm (50 mW) diode laser (240 ms exposures) with Borealis attachment (Andor), emission filter Chroma ET525/50m, and an Andor iXon3 camera. We performed imaging with a 100x 1.45 Ph3 Nikon objective and a 1.5x magnifier (built-in to the Dragonfly system). Apart from laser ablation experiments (see below), all time-lapse videos were collected with 10-second intervals. We used an Andor Micropoint attachment with galvo-controlled steering for targeted laser ablation, delivering 20-30 ns pulses of 551 nm light at 20 Hz. We imaged each cell until either spindle repair had been completed or the spindle had failed to repair after 5 min. For spindle ablation videos, we imaged every 3.5 s for control and ase1Δ videos, and every 1 s for klp6Δ and klp5Δ videos. We used Andor Fusion software to manage imaging and Andor IQ software to simultaneously manipulate the laser ablation system.

#### Live cell optical trapping and scanning confocal imaging

We performed experiments at room temperature on a LUMICKS C-trap system (LUMICKS, Netherlands). The instrument consists of a three-channel confocal fluorescence inverted microscope and dual-trap-through-the-lens optical tweezers allowing for simultaneous vesicle manipulation and confocal imaging. In the system, the sample is held between a water immersion objective (60x, NA 1.2; Nikon, Japan, MRD07602) and an oil immersion condenser (60x, NA 1.4; Leica, Germany). As such, the 1064 nm IR trapping laser beams are focused on the sample through the objective while the condenser provides brightfield illumination (850 nm LED light source). Back-scattered light from the optical trap is also detected by the photo-sensitive detectors (PSDs) and used to detect the position of the trapped object relative to the trap. To trap vesicles, we typically set the trapping laser power to 8% overall power, and adjust the split between trap 1 and 2 to 100% trap 1. We captured brightfield images on a CMOS camera with a field of view of 110 x 70 µm and simultaneously performed scanning confocal imaging with 488 nm laser line (to excite GFP) at 8% laser power, and acquired through a 525/45 emission filter on an avalanche photodiode (APD). All confocal scans had a 150 nm pixel size, resulting in a 2.15 s/frame time series. Using the brightfield images, we selected small (near diffraction-limited in size), spherical cytoplasmic structures to use as ‘handles’ for pushing on the nucleus.

It is difficult to conclusively identify these trapped ‘handles’ (Figure 2B, right panel). However, we presume they are lipid vesicles rather than lipid droplets because when we occasionally trap two of these structures simultaneously and then let go, they remain physically distinct from each other when we turn off the trap. If they were lipid droplets, we would expect them to instead fuse with each other when brought into close proximity.

### Quantification and Statistical Analysis

#### Image and video preparation and editing

To optimize the identification and tracking of spindle and nuclear envelope features, we performed initial processing of fluorescence microscopy images and videos using FIJI [42]. First, we cropped videos to spatial and temporal regions of interest and made linear adjustments to the brightness and contrast of the images, in order to track features more clearly.

For quantification of fluorescence intensity of Klp5-GFP, Ase1-GFP, and GFP-Atb2, we imaged at the same laser power and intensity and adjusted images to the same brightness and contrast settings (across all experiments). To control for mitotic stage, we performed all measurements at the spindle midzone with spindles that were 5-7 microns in length for experiments not performed on the optical trap, and collected 5 measurements per spindle at the midzone. For optical trap experiments, we measured the integrated density on the spindle 2 frames before the trap is turned on and pushing at the midzone, as well as ~40 seconds after the trap is turned off and no longer pushing. All integrated density measurements were taken with a set ROI size, spanning part of the spindle midzone. Background was measured the same way, though away from the spindle midzone. We calculated integrated density with the following expression: *Fractional IntegratedDensit* 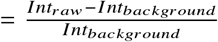. These were then averaged per cell. We tracked spindle length and curvature using home-built code, as described previously [6].

#### Tracking of spindle features in ablation videos

Following initial processing in FIJI, we performed all data quantification of post-ablation spindle dynamics via a tracking program, home-written in Python. For each ablated spindle, we tracked the two spindle poles following ablation. Using this program, and the ‘line’ tool in FIJI to measure the length of each spindle immediately prior to ablation. We then used these data to calculate the change in pole separation (length) for each spindle over time, following ablation. Additionally, we tracked the positions of the two new plus-ends of each ablated spindle half throughout the video. Our tracking program includes a method for indicating whether or not spindle repair has occurred, with the reconnection of the two ablated spindle halves, in each frame. We then used the data for frames collected before the reformation of a single spindle to compute time traces for the change in spindle half length.

We define spindle half splaying as the detectable lateral dissociation of microtubule ends from each other in ablated spindle halves. We determined splay state by eye in FIJI, using videos in which the brightness and contrast have been linearly adjusted for clarity. If a spindle half appears splayed in any frame of a video, we count that spindle half as splaying. For videos in which splay state was not readily apparent throughout, for one or both spindle halves, we made no splaying designation for the unclear half/halves.

## RESULTS

### Increased nuclear envelope tension from cerulenin treatment disrupts spindle and microtubule dynamics in *S. pombe*

We first aimed to determine how the *S. pombe* spindle dynamically responds to changing compressive force. Nearly all *S. pombe* spindle microtubule minus-ends are embedded at spindle pole bodies [10], [13], [22], making the spindle midzone a key region for both sliding-driven spindle elongation and for regulation of microtubule growth [18], [43]. When spindles buckle from increased compressive force, curvature is also concentrated in this region [6], [32]. Thus, we focused on measuring the response of the midzone region.

First, we increased nuclear envelope tension through treatment with cerulenin, a lipid synthesis inhibitor that reduces the availability of phospholipids, presumably decreasing nuclear envelope surface area and thereby increasing nuclear envelope tension. Previous work has shown this alteration in surface tension causes spindles to bend and decreases their elongation rate [6], [8], [44], [45]. Despite the changes it induces in the mitotic spindle and nuclear shape (Figure 1A), cell division still completes successfully (Figure 1B), allowing us to quantify other observable changes to the spindle.

**Figure 1.**
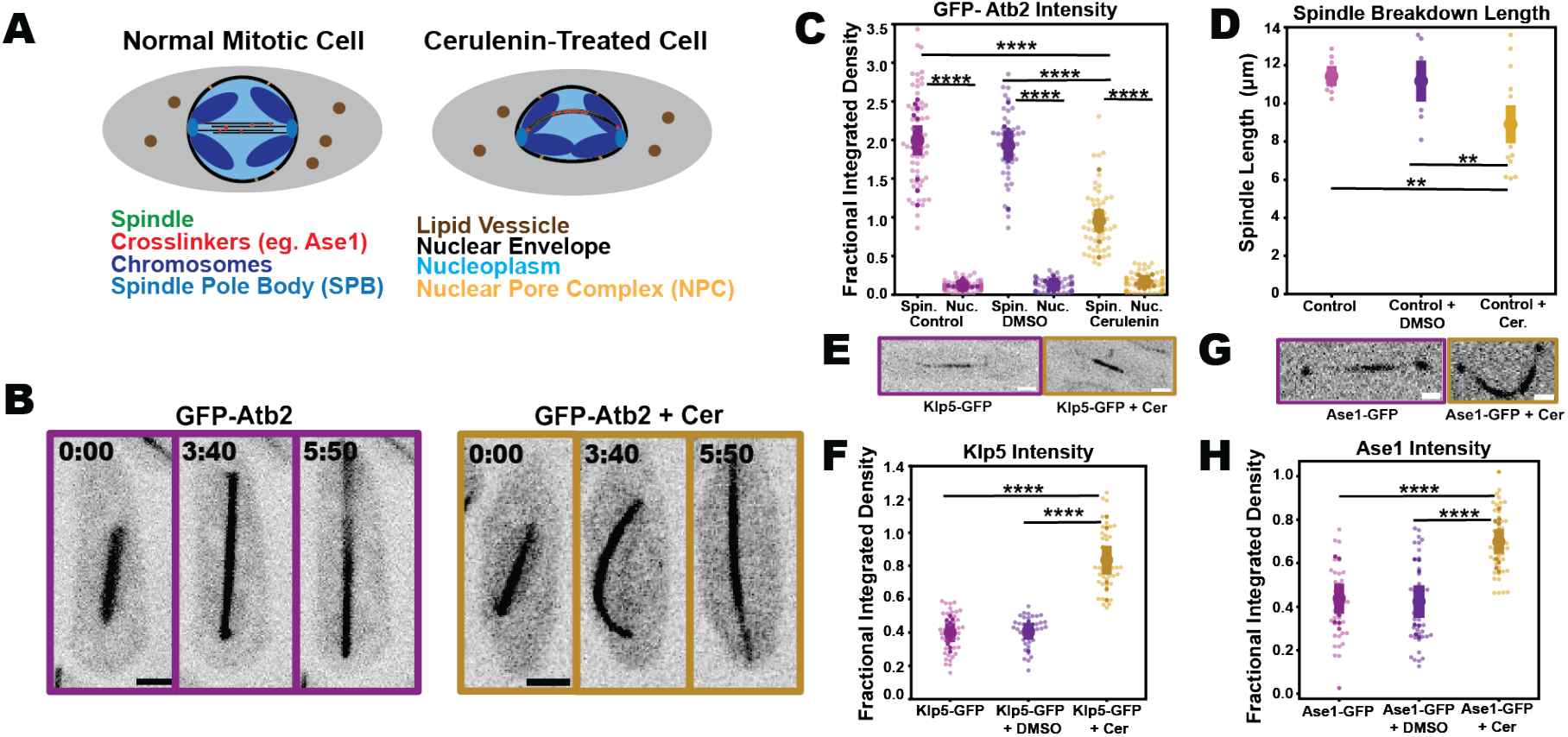
Cerulenin-induced nuclear envelope tension alters spindle regulation in *S. pombe*. Treatment conditions: 300µM cerulenin. All images are representative spinning disk confocal microscopy images; scale bars, 2 µm scale bars; time stamps, min:sec. Lighter color in all beehive plot shows all data(n), darker points indicate averages of each individual cell (N), shaded region indicates mean +/−s.e.m of cellular averages 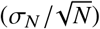. (A) Cartoon illustrates the impact of cerulenin treatment on the nuclear envelope and mitotic spindle. (B) Anaphase spindle elongation in cells expressing GFP-Atb2. Left, untreated cells; right, treated with cerulenin. (C) GFP-Atb2 fluorescence intensity in untreated, cerulenin-treated, and DMSO control cells at both the spindle midzone (Spin.) and the nucleoplasm (Nuc). (Control-Spin.: n = 60, N = 12; Control-Nuc.:n= 50, N = 10; DMSO-Spin.: n= 45, N=9; DMSO-Nuc.: n = 45, N = 9; Cer-Spin.: n=55, N = 11; Cer-Nuc.: n = 55, N = 11). (D) Spindle breakdown length in untreated (N=10), DMSO control (N=9) cerulenin-treated (N=13), and cells. (E) Representative images of untreated (left) and cerulenin-treated (right) *S. pombe* cells expressing GFP-Klp5. (F) Klp5-GFP intensity at spindle midzone (Klp5-GFP: n=45, N = 9; Klp5-GFP + DMSO: n=40, N = 8; Klp5Δ + Cer: n=45, N = 9). (G) Representative images of untreated (left) and cerulenin-treated (right) *S. pombe* cells expressing Ase1-GFP. (H) Ase1-GFP intensity at spindle midzone (Ase1-GFP: n=40, N = 8; Ase1-GFP + DMSO: n=45, N = 9; Ase1-GFP + Cer: n=50, N = 10).

**Figure 2.**
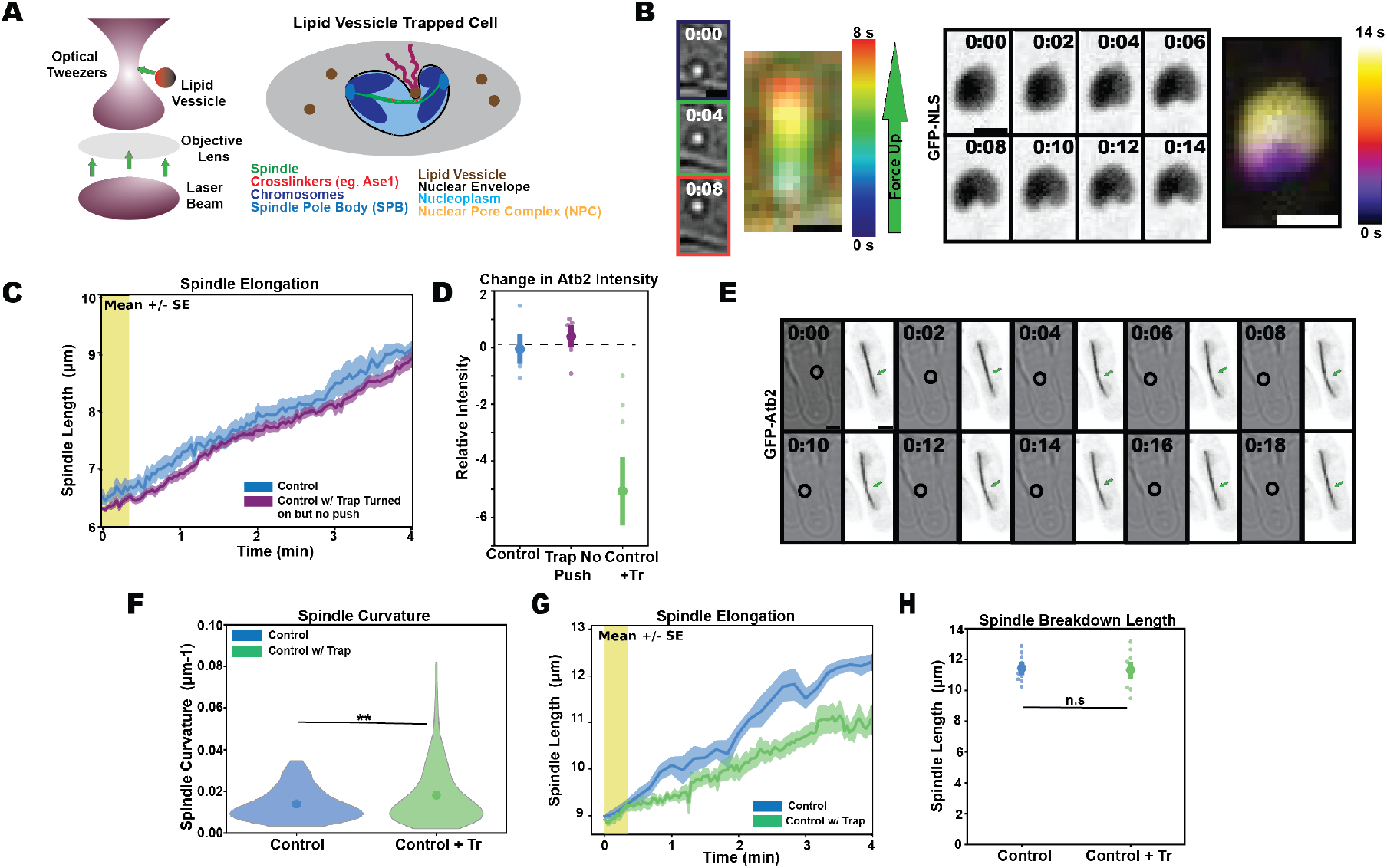
Optical trap assay applies increased force on the spindle through the nuclear envelope. (A) Assay schematic. (B) Color maps tracking lipid vesicle displacement across frames (left) with correlated time lapse of nuclear envelope deformation across (right). (C) Spindle elongation dynamics with optical trap turned off (blue, N=5) versus turned on without force exertion (purple, N=12). Traces show average spindle length, shaded region +/−s.e.m. (D) Change in integrated fluorescence intensity of GFP-Atb2 in cells imaged with optical trap off (N=6), cells imaged with optical trap on (N=5), and nuclei pushed on with the trap (N=10). (E) Representative example paired brightfield images (left) identifying the trapped vesicle (black circle) and scanning confocal images (right) identifying the pushing location (blue arrow). Scale bars, 2 µm; time stamps, min:sec. (F) Violin plots (normalized to same area) showing spindle curvature of spindles of length 5-7 µm without (N = 14, n = 163) and with (N = 10, n = 635) optical trapping. Shaded region refers to mean +/−s.e.m. Outliers (<0.44% of all data), with curvature values greater than 0.3 µm^1^ are not shown. (G) Spindle elongation showing average spindle length +/−s.e.m. in control spindles (blue, N=14) and following pushing with the optical trap (green, N=10).Yellow shaded region represents when pushing occurs. (H) Spindle breakdown length in control spindles (N=10) and spindles subjected to the optical trap (N = 8)

In these experiments, we found a reduction in the intensity of GFP-Atb2 in the midzone of the spindle, but not within the nucleoplasm (Figure 1C) suggesting that microtubule dynamics themselves, not just spindle elongation rate (which could be altered by differences in motor sliding alone), become misregulated with cerulenin treatment. Spindle breakdown length is also reduced in these cells (Figure 1D).

We next examined whether the localization of midzone components might accompany these changes to microtubules. We particularly examined Klp5, a key promoter of microtubule depolymerization, and Ase1, a crucial microtubule crosslinker. Under cerulenin treatment, we see that the intensity of Klp5-GFP along the spindle midzone approximately doubles (Figure 1E, F), while Ase1 intensity in the same region increases 1.5-fold (Figure 1G, H). The increased MAP localization to this region is particularly noteworthy given that it trends in the opposite direction to the change in microtubule intensity.

Though these cerulenin-induced changes are noteworthy, there is the possibility that cerulenin treatment might alter spindles by additional mechanisms beyond its effects on nuclear envelope membrane tension. Thus, we next sought to develop an alternative method to acutely exert compressive force on the spindle. For this approach, we turned to live cell optical trapping.

### Optically-trapped lipid vesicles can acutely exert compressive force through the nuclear envelope to bend spindles

An optical trap uses a highly focused infrared laser beam to precisely manipulate in three dimensions particles that differ in index of refraction from their surrounding medium [23]. While optically trapped lipid vesicles or granules have been previously used to push on *S. pombe* nuclei to displace them and alter their shape [23], [46], [47], the spindle response to such forces has yet to be investigated. We devised a protocol to trap lipid vesicles via optical tweezers and use the trapped vesicle as a mechanical ‘handle’ that exerts force on the spindle through the nuclear envelope barrier. By tuning the speed and timing of the force application, we induce a reproducible and consistent perturbation to the nucleus, which we use to interrogate nuclear envelope-spindle interactions and dynamics (Figure 2A).

Using brightfield imaging, we identify small cytoplasmic structures (likely lipid vesicles, see Materials and Methods) that we can successfully manipulate with the optical trap (Figure 2B). When we use these vesicles to push on the nucleus from the cytoplasm, we observe changes to nuclear shape, indicating that we can use this method to locally increase the nuclear envelope tension without disrupting its integrity (Figure 2B). Importantly, as a control, we tested whether optical trapping itself affects spindle dynamics, for example through localized heating or nonspecific photodamage. To do so, we trapped vesicles using an identical protocol but without pushing on the nucleus, and found that spindles were unaffected (Figure 2C,D).

We next tested whether acute application of force to the spindle by using the trap to push on the nuclear envelope affects spindle elongation dynamics. We pushed on spindles with sufficient force to bend them (Figure 2E), held them in the bent configuration for 10 seconds, and then released the trap and measured the ensuing response. We performed this perturbation on spindles of lengths ~5 − 7 µm, a comparable length to spindles that tended to buckle after cerulenin treatment (Figure 2E, F). Following this perturbation, spindle elongation slows, and this slowdown persists for multiple minutes after we release the force (Figure 2G), suggesting that increased inward force triggers ongoing changes to spindle length regulation, also seen in cerulenin-treated cells. We also examined effects on spindle GFP-Atb2 intensity. Because the optical trap allows spatiotemporal control of force application, we performed paired measurements of the fluorescence intensity before and after trapping, where we see a drop in intensity after exerting force with the trap (Figure 2D). We do not, however, detect a difference in spindle breakdown length following force application with the optical trap (Figure 2H), suggesting the possibility that either greater force or a longer application may be needed to recapitulate this consequence of cerulenin treatment.

Altogether, these results demonstrate that spindle microtubules are dynamically responsive to changes in force application, even when it is only applied transiently. The high correspondence between results with cerulenin and with trap-induced force - two very different approaches to changing spindle mechanics - suggests that the slowing of spindle elongation and reduction in microtubule intensity is indeed a downstream consequence of altered compressive force.

### Klp5 and Klp6 minimally contribute to regulation of spindle dynamics in response to compressive force

To better understand the underlying mechano-sensitive regulation of spindle and microtubule dynamics, we next sought to test what happens in the absence of MAPs that are typically recruited to the midzone in response to compressive force (Fig. 1E-H). We approached this question using cerulenin treatment and optical trapping: two complementary approaches that both result in increased compressive force on the spindle.

We first examined how Klp5 impacts spindle response to force application via both cerulenin and optical trapping. Overall, we find that spindles in klp5Δ cells have a similar response to changes in nuclear envelope force as control spindles without the deletion (Figure 3A). Compressed spindles still exhibit reduced microtubule intensity, equivalent to that seen in treated control cells (Figure 3B). These spindles also curve to a similar extent to control cells (Figure 3C), suggesting overall analogous mechanical properties and responsiveness. While klp5Δ alone reduces spindle elongation rate [2], we see only a small additional decline in elongation rate under cerulenin treatment (Figure 3D) and no further change in that rate after pushing on spindles with the optical trap (Figure 3E). We find, as has previously been measured, that klp5Δ results in slightly extended lengths at spindle breakdown, likely due to a reduction in microtubule depolymerization [2]. In a qualitatively similar response to force as control cells, this length is slightly reduced in both klp5Δ cells treated with cerulenin and, to a smaller extent, in nuclei subjected to pushing forces from the optical trap (Figure 3F). Altogether, these results suggest that klp5Δ only slightly attenuates the response of the spindle to compressive force. We thus conclude that it is unlikely to be the driving regulator of force-dependent changes in the spindle microtubule apparatus, and is also not crucial for maintaining force balance between the nuclear envelope and the spindle.

**Figure 3.**
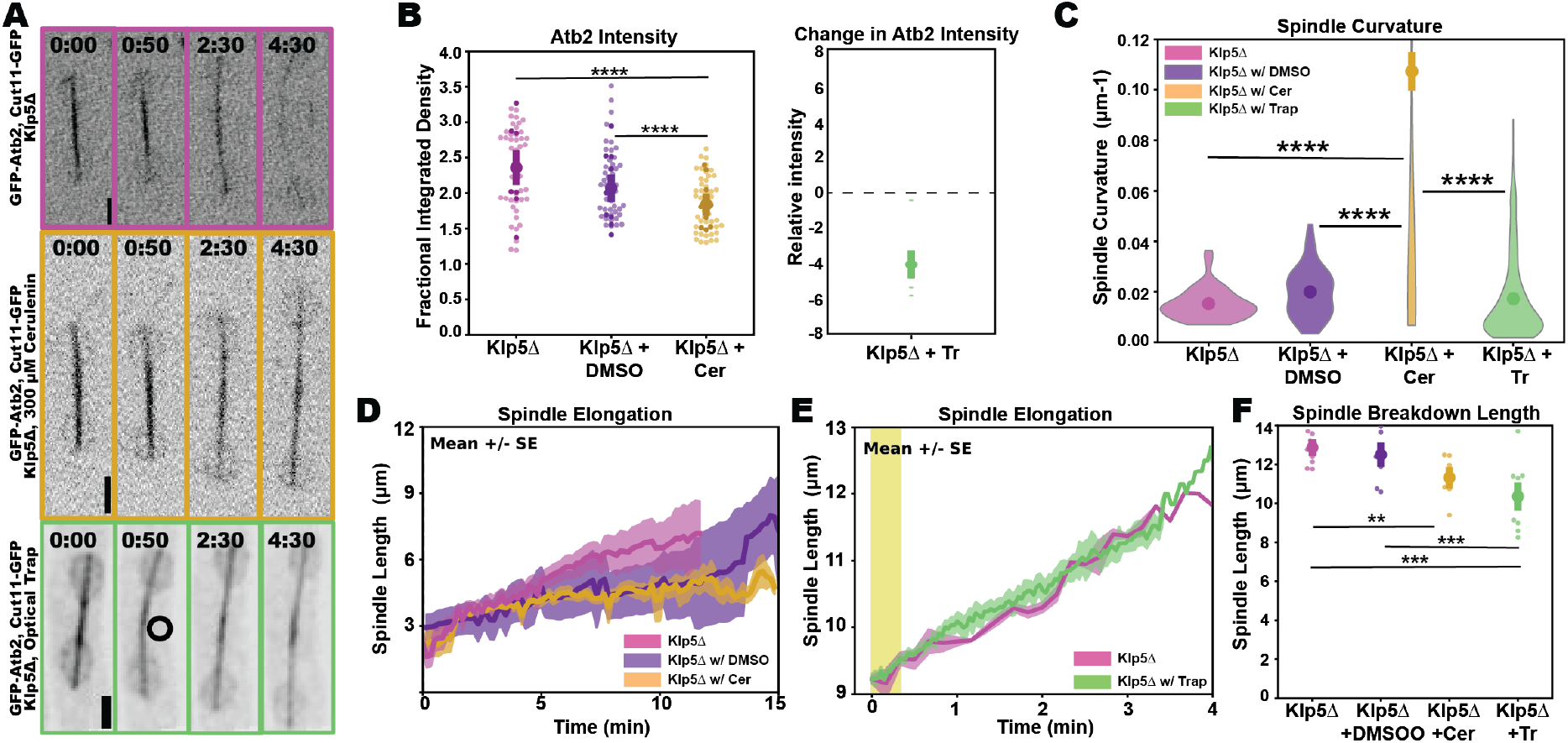
Loss of Klp5 minimally affects spindle response to compressive force (A) Representative time-lapse panel of spindle elongation in klp5Δ *S. pombe* expressing GFP-Atb2 and GFP-Cut11 (MWE75) with no treatment (top), cerulenin treatment (middle) and optical trapping (bottom, circle indicating trapped veiscle). Scale bars, 2 µm; timestamps, min:sec. (B) Left: Fluorescence intensity (of GFP-Atb2) with klp5Δ in untreated (N = 11, n = 55), DMSO control (N = 11, n= 55), and cerulenin-treated (N = 10, n = 50) cells. Lighter color in beehive plot shows all data (n), darker points indicate averages of each individual cell (N), shaded region indicates mean +/−s.e.m.. Right: Average change in fluorescence intensity in klp5Δ spindles subjected to the optical trap (as in Figure 2C) (N = 10). (C) Violin plots showing spindle curvature of klp5Δ cells at 5-7 µm of length under no treatment (magenta, N = 13, n = 130), DMSO treatment (purple, N = 9, n = 124), cerulenin treatment (gold, N = 16, n = 235), and optical trapping (light green, N = 10, n = 828). The area of each violin is normalized to the same area. Shaded region highlights the mean +/−s.e.m. Outliers (< 0.62% of all data), with curvature values greater than 0.3 µm^1^ are not shown here. (D) Spindle elongation in klp5Δ (magenta, N = 13), klp5Δ with DMSO (purple, N = 9) and klp5Δ with cerulenin treatment (gold, N = 16). Traces represent the average spindle arc length while shaded regions represent average +/−s.e.m. Traces are aligned such that t=0 is the time when the rate of spindle elongation sharply increases. (E) Spindle elongation in untrapped (magenta), N = 10 versus trapped (light green, N = 10) klp5Δ spindles. Traces are aligned such that t=0 is the length when lipid vesicles begin pushing against the spindle. The yellow shaded region highlights when the trap is turned on. (F) Spindle breakdown length in untreated klp5Δ (magenta, N = 10), DMSO-treated klp5Δ (purple, N = 10) cerulenin-treated klp5Δ (gold, N = 10), and optical trap-treated klp5Δ (light green, N = 10).

Due to its role as a partner of Klp5, we also asked whether Klp6 contributes to spindle mechanosensitivity. Upon compressing the spindle via both cerulenin treatment and the optical trap, klp6Δ spindles show a decrease in microtubule intensity (Figure 4A, B), though this effect does seem somewhat reduced compared to control spindles. Spindles deficient in Klp6 additionally curve with both cerulenin and the optical trap, illustrating that they are mechanically similar to treated control spindles (Figure 4A, C). Like klp5Δ, klp6Δ alone reduces spindle elongation rate and causes a slight increase in spindle breakdown length [2]. In this case, force exertion does not seem to cause additional impact on spindle elongation rate or spindle breakdown length, indicating that klp6Δ spindles are somewhat less mechanically sensitive than control spindles (Figure 4D-F). Overall, while spindles in klp6Δ remain responsive to force in ways that are qualitatively similar to control cells, klp6Δ appears to somewhat decrease the sensitivity of the response. We thus conclude that, while Klp6 may make some contribution to the spindle’s response to force, like Klp5, it is not essential for these effects and is dispensable for maintaining the spindle-nuclear envelope force balance.

**Figure 4.**
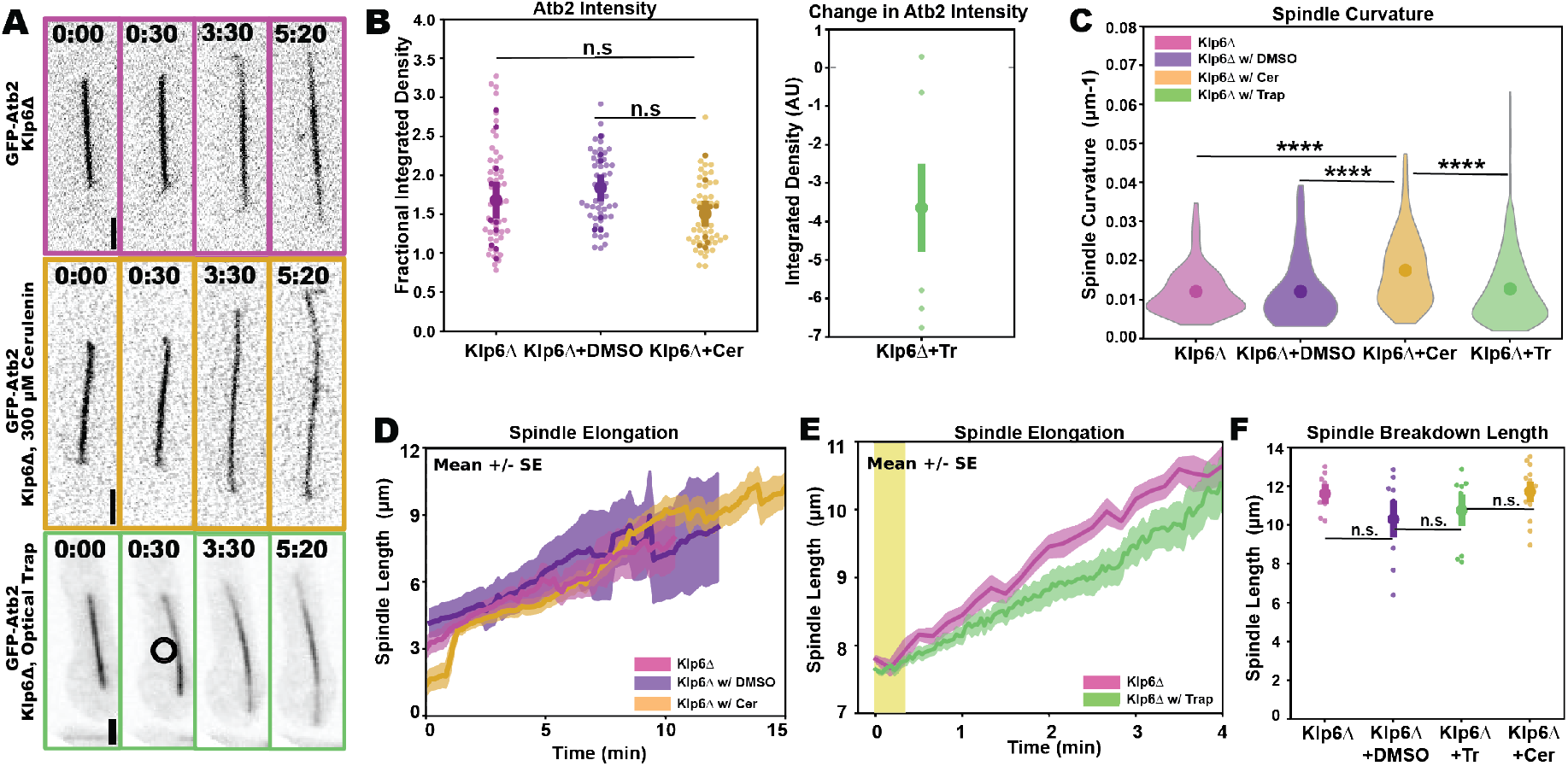
Loss of Klp6 minimally affects spindle response to compressive force. (A) Time-lapse panel of spindle elongation in klp6Δ *S. pombe* expressing GFP-Atb2 (MWE76) with no treatment (top), cerulenin treatment (middle) and optical trapping (bottom, circle indicates trapped vesicle). Scale bars, 2 µm; timestamps, min:sec. (B) Left: Fluorescence intensity (of GFP-Atb2) in untreated (N = 11, n = 55), DMSO control (N = 10, n = 50), and cerulenin-treated (N = 10, n = 50) cells. Lighter color in beehive plot shows all data points (n) darker points indicate averages of each individual cell (N), shaded region indicates mean +/−s.e.m of cellular averages 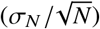 Right: Average change in fluorescence intensity in klp6Δ spindles subjected to the optical trap (as in Figure 3C) (N = 8). (C) Violin plots showing spindle curvature of klp6Δ cells at 5-7 µm in length under no treatment (magenta, N = 12, n = 223), DMSO control (purple, N = 11, n = 157), cerulenin treatment (gold, N = 19, n = 330), and optical trapping (light green, N = 10, n = 1006). Violins are normalized to the same area. Shaded region highlights the mean +/−s.e.m. Outliers (< 0.57% of all data), with curvature values greater than 0.3 µm^1^ are not shown. (D) Spindle elongation in klp6Δ (magenta, N = 12), klp6Δ with DMSO (purple, N = 11) and klp6Δ with cerulenin treatment (gold, N = 19). Traces represent the average spindle arc length while shaded regions represent average +/−s.e.m. Traces are aligned such that t=0 is the time when the rate of spindle elongation sharply increases. (E) Spindle elongation in untrapped (magenta, N = 12) versus trapped (light green, N = 10) klp6Δ spindles. Traces represent average spindle arc length and shaded regions represent average +/−s.e.m. Traces are aligned such that t=0 is the length at which lipid vesicles push against the spindle. The yellow shaded region highlights when the trap is turned on. (F) Spindle breakdown length in untreated klp6Δ (magenta, N = 9), DMSO control klp6Δ (purple, N = 9), cerulenin-treated klp6Δ (gold, N = 15), and optical trap-treated klp6Δ (light green, N = 10).

### Increased nuclear compressive force rescues spindle dynamics and elongation in the absence of Ase1

Finally, we tested the responsiveness of cells with ase1Δ to nuclear compressive forces. These spindles responded strikingly differently to compressive force than control spindles (Figure 5A), especially compared to the effects of ase1Δ alone. Previous work shows that, with ase1Δ, spindles are noticeably less stable, exhibiting a phenotype where they often grow and collapse as they elongate [18]. In contrast, ase1Δ cells respond to compressive force, induced either by cerulenin or by the optical trap, by a notable increase in GFP-Atb2 intensity along the spindle (Figure 5B). In other words, in the absence of Ase1, the spindle responds to increased compressive force by stabilizing microtubules. Spindles of ase1Δ cells are also less likely to curve in response to compressive force (Figure 5C), a change that may result from an increase in spindle stiffness, potentially due to an increased number of spindle microtubules. It might also be that, without crosslinking in the midzone by Ase1, spindle microtubules may sometimes slide past each other or fray apart rather than bend. Indeed, we sometimes see splaying instead of bending under compression in these cells (Figure 5A, bottom panel). Spindles in ase1Δ cells treated with cerulenin (Figure 5D) or compressed with the optical trap (Figure 5E) both elongate at a rate that approaches that of control spindles, and are considerably faster than ase1Δ spindles alone. This is especially the case in spindles subjected to optical trapping forces, which tend to stall in elongation. Thus, ase1Δ and compressive force together actually combine to rescue each phenotype compared to either perturbation alone. Further, ase1Δ spindles subjected to force from the optical trap breakdown at slightly shorter lengths despite their increased elongation speed, while cerulenin-treated ase1Δ cells breakdown at roughly normal lengths (Figure 5F). We thus conclude that the Ase1 recruited to the spindle midzone in response to compressive force is likely required for the down-regulation of microtubules that accompanies spindle compression. In contrast, in the absence of Ase1, spindles respond to increased compressive force by stabilizing microtubules and resuming normal elongation speed.

**Figure 5.**
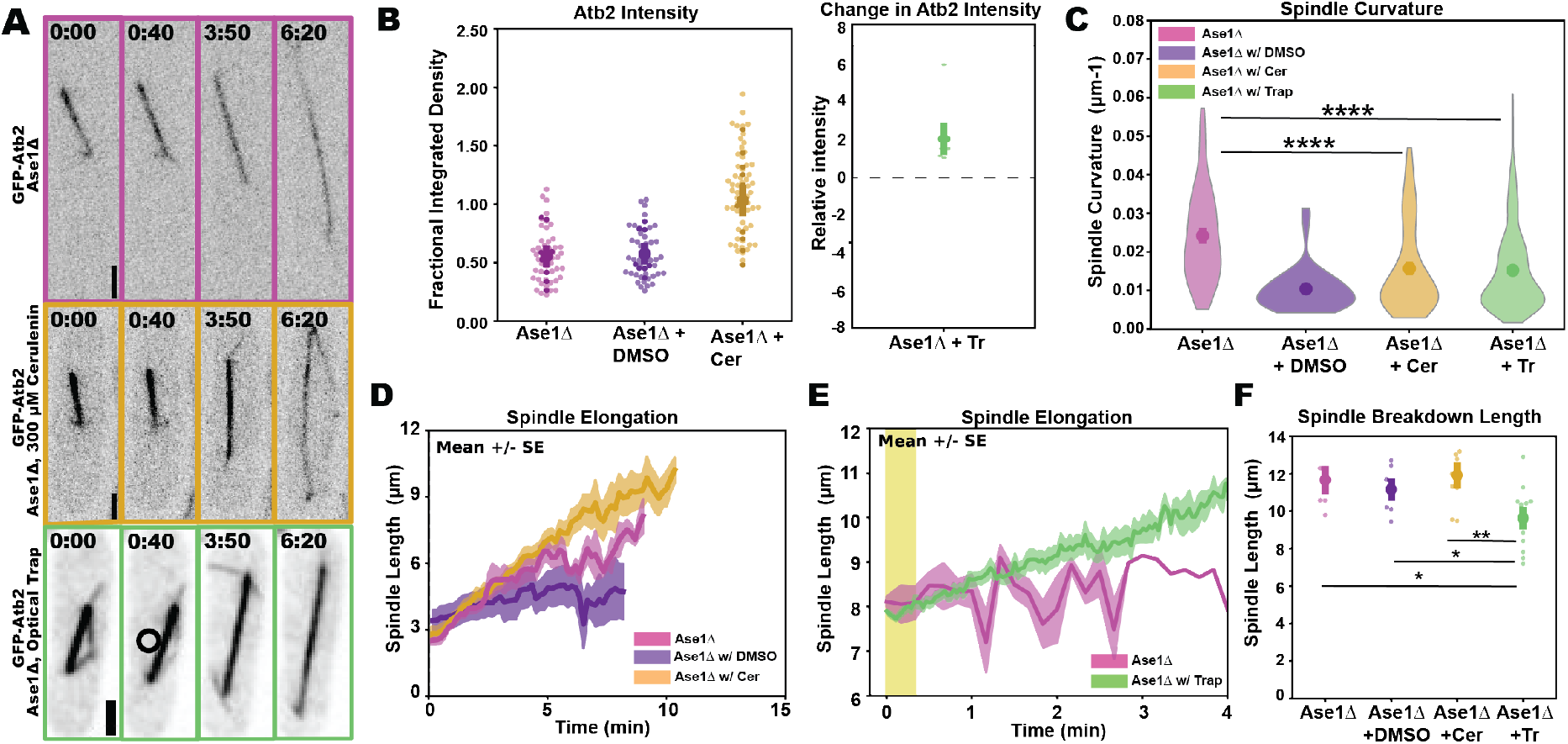
While Ase1 is crucial for spindle stability, compressive force rescues spindle and microtubule dynamics in its absence. (A) Representative time-lapse panel of spindle elongation in ase1Δ *S. pombe* expressing GFP-Atb2 (MWE3) with no treatment (top), cerulenin treatment (middle) and optical trapping (bottom, circle indicates trapped vesicle). Scale bars, 2 µm; timestamps, min:sec. (B) Left: Fluorescence intensity (of GFP-Atb2) in untreated (N = 10, n = 50), DMSO control (N = 9, n = 45, and cerulenin-treated (N= 13, n = 65) cells. Lighter color in beehive plot shows all data (n), darker points indicate averages of each individual cell (N), shaded region indicates mean +/−s.e.m of cellular averages 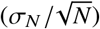. Right: Average change in fluorescence intensity in ase1Δ spindles subjected to the optical trap (N = 7). (C) Violin plots showing spindle curvature of ase1Δ cells at 5-7 µm in length under no treatment (magenta, N = 8, n = 118), DMSO treatment (purple, N = 11, n = 84), cerulenin treatment (gold, N = 10, n = 190), and optical trapping (light green, N = 14, n = 1217). Violin are normalized to the same area. Shaded region highlights mean +/−s.e.m. Outliers (<0.57% of all data), with curvature values greater than 0.3 µm^1^ are not shown. (D) Spindle elongation in ase1Δ (magenta, N = 8), ase1Δ with DMSO (purple, N = 11) and ase1Δ with cerulenin treatment (gold, N = 10). Traces represent the average spindle length +/−s.e.m. (shaded area). Traces are aligned such that t=0 is the time when the rate of spindle elongation sharply increases. (E) Spindle elongation in untrapped (magenta, N = 8) versus trapped (light green, N = 14) ase1Δ spindles. Traces represent average spindle length +/−s.e.m. (shaded region). Traces are aligned such that t=0 is the length at which lipid vesicles push against the spindle. The yellow shaded region highlights when the trap is turned on. (F) Spindle breakdown length in untreated ase1Δ (magenta, N = 8), DMSO-treated ase1Δ (purple, N = 9), cerulenin-treated ase1Δ (gold, N = 10), and optical trap-treated ase1Δ (light green, N = 12).

### Ase1 slows microtubule depolymerization in response to laser ablation

After finding that the absence of Ase1 stabilizes spindle dynamics in the presence of increased compressive force from the nuclear envelope, we sought to further elucidate how Ase1 may affect microtubule dynamics. To assess microtubule stability in the presence and absence of Ase1, we severed spindle microtubules in the spindle midzone via laser ablation. Previous work in other biological systems has shown that most severed microtubules respond to laser ablation by undergoing rapid catastrophe (depolymerization) from the plus-end, due to the removal of the stabilizing GTP ‘cap’ normally present on growing microtubule plus ends [48], [49]. In contrast, our previous work showed that severed *S. pombe* spindles are unusually stable following severing by laser ablation [25] (Fig. 6A).

**Figure 6.**
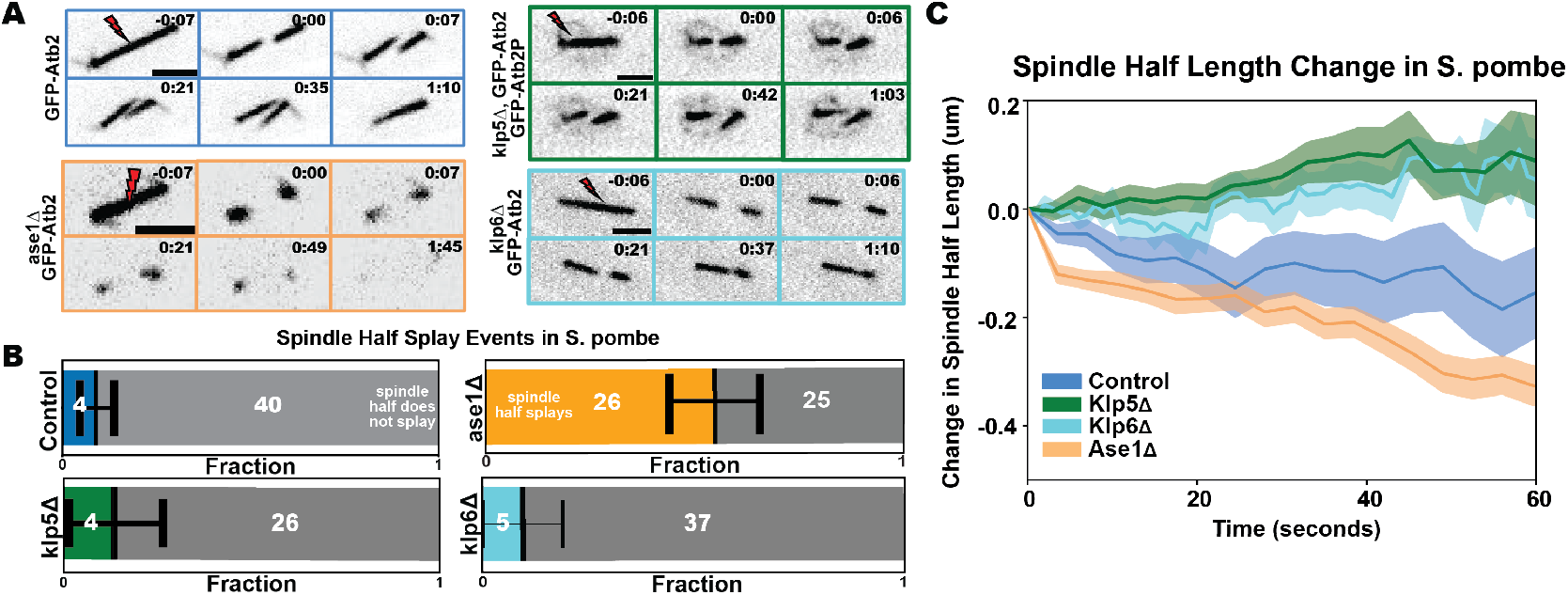
Ase1 prevents depolymerization and stabilizes microtubules in response to laser ablation (A) Time-lapse panel of spindle repair response to ablation in cells expressing GFP-Atb2. Control (blue), ase1Δ (orange), klp6Δ (cyan) and klp5Δ (green). Scale bars, 2 µm and timestamps, min:sec. (B) Counts showing splay events for control, ase1Δ klp5Δ, and klp6Δ *S. pombe*, with Ase1 displaying significantly more splaying events. (C) The change in spindle half length in control (blue, N = 13, n = 26), klp5Δ (green, N = 11, n = 21), ase1Δ (orange, N = 13, n = 20), and klp6Δ (cyan, N = 26, n = 52). Traces represent the average change in spindle half length +/−s.e.m. (shaded region). Traces are aligned such that t=0 is the frame after ablation is complete.

In striking contrast to control spindles, ablated spindles of ase1Δ cells rapidly depolymerize over the course of roughly 2 minutes (Figure 6B, E). Consistent with Ase1’s known crosslinking role, microtubules in these spindles also tend to splay apart from each other much more often than control spindles (Fig. 6F). Spindles of klp5Δ and klp6Δ cells behave more like control spindles in response to ablation. (Fig. 6C, D).

Together, these data suggest that, in addition to its known role as a spindle crosslinker, Ase1 has an additional function in regulating microtubule dynamics.

## DISCUSSION

Here, we show that microtubule regulation responds dynamically to nuclear force exerted on the spindle in closed mitosis of *S. pombe*. We alter nuclear envelope tension using two distinct approaches: the pharmacological treatment of cells with cerulenin and the direct application of force on nuclei by an optically trapped lipid vesicle. These two distinct approaches yield highly congruent changes in the mitotic spindle (Figure 7). In both cases, increased nuclear envelope tension is accompanied by decreased microtubule density and slowed spindle elongation. While cerulenin treatment causes recruitment of molecular motor Klp5 and microtubule crosslinker Ase1 to the spindle midzone, changes in microtubule regulation are dependent on Ase1, but not on Klp5 and Klp6. Additionally, Ase1 prevents microtubule depolymerization in response to laser ablation. These results point toward a previously unknown feedback loop between force, spindle protein affinity, and spindle microtubule dynamics.

**Figure 7.**
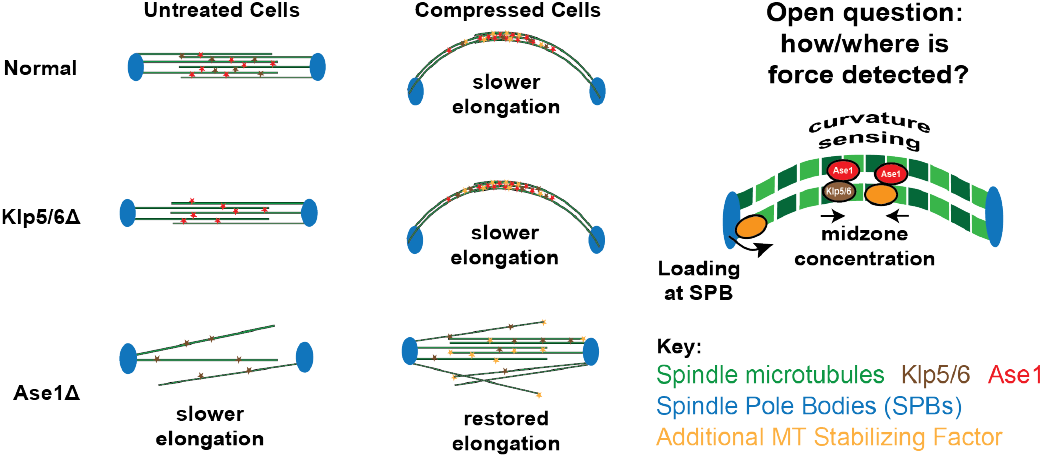
In normal cells, compressive force reduces spindle bending and causes a reduction in microtubule intensity and increased recruitment of Ase1 and Klp5 to the midzone. In klp5Δ and klp6Δ cells, the mechanical response is unchanged relative to control spindles. In contrast, with ase1Δ, compressive force induces an increase in microtubule intensity. The mechanism by which force triggers this biochemical response remains an open question for future research.

Decades of important effort have helped to elucidate the role of crosslinkers and motor proteins in regulating microtubule dynamics and spindle elongation in *S. pombe* [2], [14], [15], [17], [18], [25], [33], [34], [37], [50]. However, because of the challenge of altering force directly, it has remained difficult to disentangle biochemical and mechanical aspects of spindle signaling from each other. Our use in this study of two very disparate methods - one pharmacological and one physical - with such qualitatively consistent responses strongly suggests an upstream role for mechanical force in spindle regulation.

While the results of this study support that mechanical force causes downstream biochemical changes in the spindle, how this force is ‘read out’ by spindle proteins, including Klp5 and Ase1, remains an important question (Figure 7). We can see three potential alternatives, each of which is particularly well-suited for future *in vitro* studies that might best distinguish between them. First, we note that spindle curvature is a notable consequence of increased compressive force, and that this geometry could be detected by spindle proteins, which could be preferentially recruited to curved microtubules. Secondly, spindle proteins might be concentrated by forces between microtubules. There is already some *in vitro* evidence that such a mechanism occurs in microtubule crosslinkers, including budding yeast homologs of Ase1, which can become concentrated in regions of overlap due to microtubule sliding [51]. Furthermore, some microtubule-binding proteins can become concentrated due to preferential sliding directions [52]. Finally, increased compressive force on the spindle might be detected at the spindle pole body, where this additional force is actually exerted, and motors or other MAPs could then load additional proteins onto the spindle from there. Of course, it is possible that compressive force detection includes a multitude of these mechanisms rather than only one.

A second open question is how initial detection of compressive force is fed forward to regulate both microtubule number and spindle elongation. One possibility is that accumulated microtubule-associated factors that detect compressive force directly alter microtubule dynamics and/or spindle sliding speed. A second possibility is that changes in protein density and/or microtubule sliding speed might have downstream effects that cause further changes in the accumulation of other spindle factors. However, once one considers even a few such factors, complexity increases rapidly. Thus, computational modeling is likely to be a fruitful approach that allows the integration and assessment of these varied hypotheses.

Finally, the molecular details of Ase1’s involvement in this regulation are a particularly interesting question. Since Ase1 crosslinks microtubules in the spindle, we initially expected it to help stabilize and potentially to increase microtubule density, either directly or through its association with the microtubule stabilizer Cls1 [19]. Instead, accumulation of Ase1 in the spindle midzone in response to compressive force seems associated with fewer microtubules and slowed elongation. How does this occur? Slowed elongation is somewhat easier to explain: Ase1 has been demonstrated to serve as a ‘brake’ on spindle elongation in budding yeast [15], essentially making a frictional contribution that slows spindle elongation, and may do the same thing in *S. pombe*. To explain the reduction in microtubule density, one hypothesis is that perhaps Ase1 binds so densely to the spindle midzone that it competes off other microtubule binders that have a greater capacity for bundling or elongation, such as the molecular motor Klp5. When Ase1 is deleted, these other factors may more readily bind to this region. Under this model, Ase1 does not stabilize the spindle under force, but rather restrains the recruitment of factors that would otherwise do so.

## CONCLUSION

Overall, the analogous spindle responses we see from both cerulenin treatment and optical trapping suggest that force changes on the nuclear envelope can regulate the spindle regardless of whether force is induced globally or locally. Furthermore, we demonstrate that optically trapping specifically grants us insight into spindle stability. Through direct and controlled force exertion, we illustrate how cross-linkers like Ase1 are crucial for spindle stability and elongation dynamics, whereas motor proteins like Klp5 and Klp6 are required for microtubule depolymerization, but dispensable for bundle stabilization. Finally, we reveal that forces exerted from the nuclear envelope can act as a mechanical cue to reinforce spindle stability by promoting increased microtubule density. In the future, a deeper understanding of this mechanistic response to compressive force would allow us to further analyze this feedback loop, and perhaps identify other spindle stabilizing factors.

## AUTHOR CONTRIBUTIONS

**Experimental design:** TM, MB, MWE, CN, RH **Experimental data collection and analysis:** TM, MB, CPM, MWE, CN. **Manuscript writing and editing:** TM, RH, MWE. **Obtaining funding:** MWE. All authors have given final approval of the submitted version.

## ACKNOWLEDGMENTS

We thank M. Betterton and members of the Elting lab, for advice and helpful discussions, and we thank F. Chang, M. Betterton, J. R. McIntosh and C. Laplante for strains. We thank Nataional Bio-Resource Project - Yeast,Japan (NBRP) for strain MWE60 - FY14688. We thank the Weninger Lab (NCSU) and Wang Lab (NCSU) for sharing lab space and equipment. We thank the Cellular and Molecular Imaging Facility (CMIF) at NCSU, which is supported by the State of North Carolina and the National Science Foundation, for microscopy support. We thank the Light Microscopy Core Facility (LMCF) at Duke University for imaging and optical trap support, supported by NIH 1S10OD30407-01A1. This work was supported by NIH 1R35GM138083 and NSF 2133276.

